# Modelling ferroptosis in a human microglial line by sequential exposure to iron and GPX4 inhibition

**DOI:** 10.64898/2026.01.19.700282

**Authors:** Renaud Bussiere, Nikhil Tulsian, Cecilia Wieder, Dewi McConnaughie, Evie Tynan, Andrew Lowe, Esther Cheow, Matthew Choo, Jill Richardson, James A. Duce, Sébastien Gillotin

## Abstract

Excessive iron accumulation is a pathological feature of several neurodegenerative diseases (NDDs) and a growing body of evidence suggests that ferroptosis, an iron-dependent form of regulated cell death (RCD) driven by lipid peroxidation, is implicated in their pathogenesis. Microglia, the brain’s resident immune cells, buffer iron overload but become susceptible to ferroptotic death, exacerbating neuroinflammation and neuronal loss. To uncover the molecular events leading to microglial ferroptosis, we established a human microglial ferroptosis model using the HMC3 cell line. This model recapitulates core features of ferroptosis, including increased reactive oxygen species (ROS) and peroxidation of lipids at the membrane, both rescued by Ferrostatin-1 (Fer-1). We used this model to perform integrated multi-omics profiling and identified significant dysregulation in lipid species, notably an accumulation of sterols, including oxysterols such as the 7-oxo-cholesterol, alongside the oxidation of polyunsaturated fatty acid (PUFA) characteristic of ferroptosis. Transcriptomic and proteomic analyses corroborated these findings, revealing the upregulation of genes and proteins involved in the mevalonate pathway and cholesterol metabolism. Importantly, the increased expression of some of these key metabolic genes was also reversed by Fer-1 treatment, indicating their role in a pre-ferroptotic signature. Our model provides a novel platform for investigating early molecular events in microglia ferroptosis. Integrating these findings into future investigations could uncover new protective mechanisms against microglia ferroptosis at the crossroad between ROS level mitigation and sterol metabolism.

## Introduction

Ferroptosis is a non-apoptotic, iron-dependent form of regulated cell death (RCD) defined by the accumulation of lipid peroxides in cellular membranes^1,2^. The cystine transporter, xCT, and the glutathione peroxidase 4 (GPX4) are key for antioxidant cellular defence to detoxify lipid peroxides and prevent ferroptosis^2–5^. Intracellularly, organelles involved in lipid metabolism as well as reactive oxygen species (ROS) production and consumption, such as the endoplasmic reticulum (ER), peroxisomes and mitochondria emerge as hotspots for ferroptosis initiation^6–9^.

While polyunsaturated fatty acids (PUFAs) are generally viewed as the primary substrates for lipid peroxidation^10^, the overall lipid composition of cellular membranes is a critical determinant of ferroptotic sensitivity. Other lipid classes have been implicated in ferroptosis regulation. On one hand, ether phospholipids that contribute to the stability and fluidity of the membrane^11^ promote ferroptosis when oxidised^8,12^, whilst sterols including cholesterol can form a structural component of membranes synthesised via the mevalonate pathway to increase ferroptosis resistance^13,14^. This resistance by the mevalonate pathway is strengthened through the involvement of components, such as GPX4, squalene and CoQ10^15^, as well as other intermediates acting as radical trapping agents. Indeed, inhibition of the cholesterol synthesis enzyme DHCR7 has recently been demonstrated as a mechanism to promote survival against ferroptosis via accumulation of 7-dehydrocholesterol (7-DHC)^16,17^. A key consideration in ferroptotic sensitivity is that the lipid composition affected by oxidation is likely to be cell type specific and thus understanding which lipid classes mostly contribute to a ferroptotic phenotype in microglia is of high interest but yet to be fully elucidated.

Ferroptosis is implicated in multiple physiological^18^ and pathological processes, such as tissue damage associated with ischemia/reperfusion^19^, liver diseases^20^, acute renal failure^21^ and could contribute to both oncogenic and tumour suppressor properties^22–24^. In the brain, iron accumulation is thought to be associated with pathological hallmarks of several neurodegenerative diseases (NDDs)^25–27^, including Parkinson’s Disease (PD)^28,29^, Huntington’s Disease (HD)^30^, Amyotrophic Lateral Sclerosis (ALS)^31^ and Alzheimer’s Disease (AD)^32^. Microglia have higher iron-storage capacity than neurons^33^, but in NDDs their uptake of iron and phagocytose iron-rich debris^34,35^ results in an increase burden on the intracellular anti-oxidant and inflammatory defensive mechanisms. Moreover, microglial susceptibility to ferroptosis may vary depending on their activation phenotype^36,37^ and recent evidence of non-cell autonomous neuronal death was associated with sublethal ferroptotic stress ^38^. The release of large quantities of intracellularly accumulated iron into the extracellular environment is a major consequence of microglial ferroptotic death that contributes to fuelling a negative environment for surrounding neurons. Furthermore, ferroptotic microglia increase pathways involved in vesicle trafficking which concurs with neurodegeneration^39^. Additional efforts in identifying gene signatures protecting or sensitising microglia to ferroptosis would support better approaches to tackle mechanisms of neurodegeneration associated with ferroptosis.

To support outstanding questions in microglia ferroptosis, we endeavoured to engineer a reproducible model of human microglia ferroptosis using the HMC3 cell line. We specifically designed a sublethal ferroptotic phenotype by loading HMC3 with iron followed by an acute dose of RSL3 that is suitable to investigate early ferroptotic events. Using this model, integrated multi-omics profiling (transcriptomics, proteomics, lipidomics) identified cholesterol metabolic pathways as an initial defensive mechanism in iron-laden cells. The inhibition of GPX4 with RSL3 in these cells increases ROS levels and oxysterols to heighten sensitivity to ferroptosis. Altogether, our work provides a straightforward platform for further investigations of pre-ferroptotic pathways in microglia and paves the way for identifying targets that could be evaluated in microglia models of NDDs.

## Results

### Iron-loading primes RSL3-induced ferroptosis in human HMC3 microglial cell line

To interrogate signalling pathways involved in the triggering of ferroptosis, HMC3 were used to engineer a sublethal microglial model that recapitulates molecular features of ferroptosis. As a key regulator of ferroptosis through blockage of Glutathione Peroxidase 4 (GPX4), the inhibitor RSL3^40^ at a range of concentrations (10 nM to 1 µM) was acutely dosed (2 hr) to evaluate lipid peroxidation induction and cell survival (Fig. 1A). Lipid peroxidation increased in a dose dependent manner (Fig. 1A i) to reach a plateau at the top concentration (i.e. 1 µM), but more than 80% of the cells still survived at this concentration (Fig. 1A-ii), suggesting that despite the lipid peroxidation induction, the timeframe was not sufficient to cause the cell death observed at later time points (Sup Fig. 1). In contrast, a less potent activator of ferroptosis Erastin (IC_50_ of 10 µM^1^) did not induce lipid peroxidation in the same timeframe (Fig. 1B) supporting the use of RSL3 as the preferred inducer of lipid peroxidation in this microglial model.

**Figure 1:**
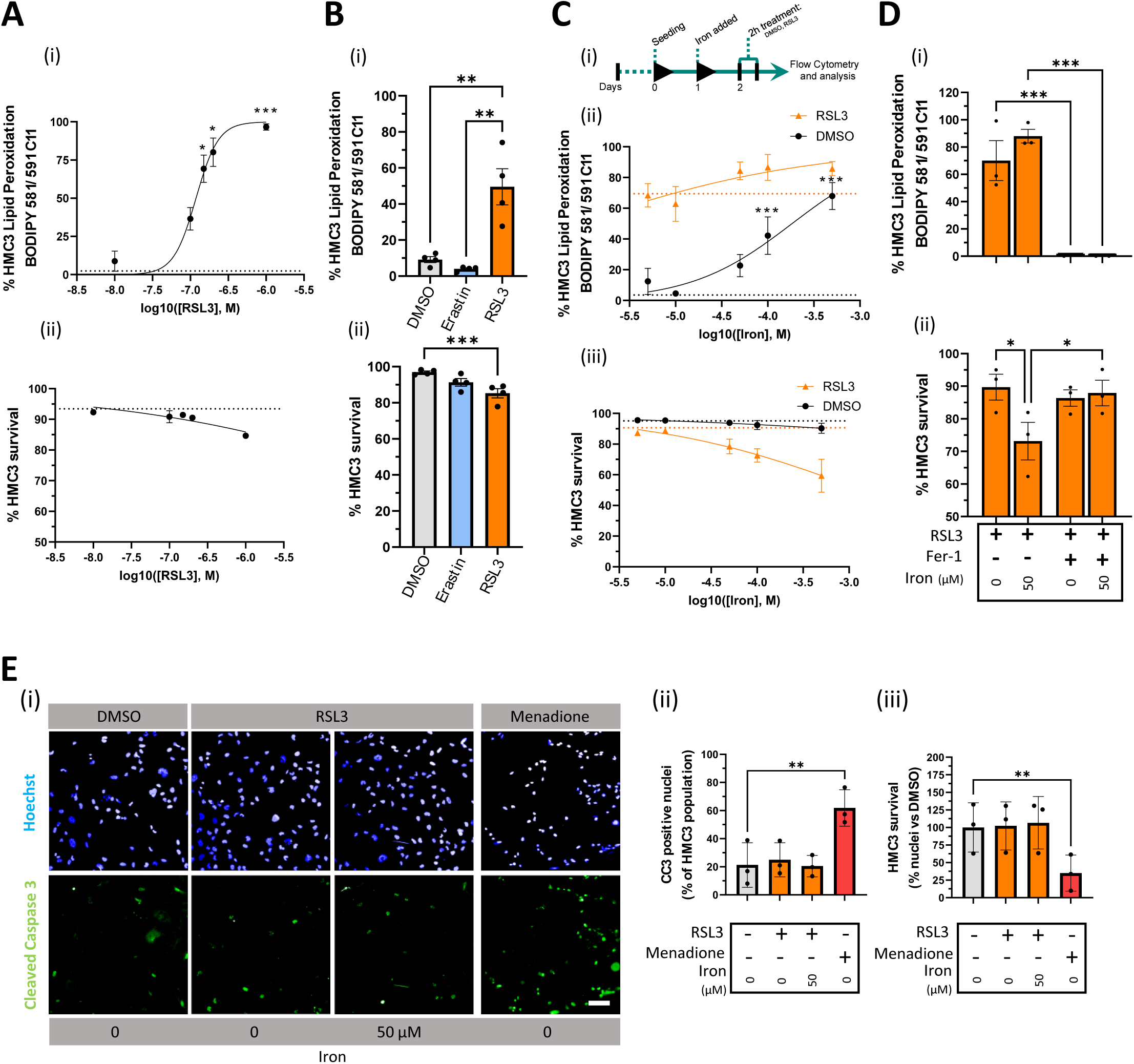
Ferroptosis modelling in HMC3 microglia. A) Dose-response curves with RSL3 treatment (0-1 µM) for 2 hr with measurement of (i) levels of lipid peroxidation (BODIPY 581/591 C11) and (ii) cell viability (DRAQ7). Dotted line represents control values from DMSO-treated samples against which other samples are compared for statistical significance. N=3, one-way ANOVA with Šidák’s test, *p < 0.05, ***p < 0.001. B) Cytofluorimetric measurement of (i) levels of lipid peroxidation and of (ii) cell viability, following treatment with Erastin (10 µM), RSL3 (200 nM) or DMSO (control). N=3, one-way ANOVA with Tukey’s test, **p < 0.005, ***p < 0.001. C) (i) Overview of the experimental design. Dose-response curves of iron-loading (0-500 µM) for 24 hr, followed by treatment with DMSO (circles) or RSL3 (200 nM, triangles) for 2 hr measuring levels of (ii) lipid peroxidation and (iii) cell viability. Dotted lines represent control values from samples without iron against which the other samples are compared for statistical significance. N=3, one-way ANOVA with Dunett’s test, *p < 0.05. D) Rescue phenotype with Fer-1 (10 µM) after single or combined treatments of iron (50 µM) and RSL3 (200 nM) measuring levels of (i) lipid peroxidation and (i) cell viability. N=3, one-way ANOVA with Tukey’s test, *p < 0.05, ***p < 0.001. E) Representative images of HMC3 cells stained with (i) CC3 antibody (Green). Menadione treatment was used as positive control to induce apoptosis. (ii) Quantification of CC3 positive nuclei, plotted as percentage of total number of nuclei defined with Hoechst. Scale bar represents 100 µm. (iii) HMC3 survival measuring total nuclei (Hoechst) and plotted as percentage over DMSO-treated control. N=3, one-way ANOVA with Tukey’s test, **p < 0.01.

To recapitulate the iron-enriched microenvironment associated with neurodegenerative disease, ferric ammonium citrate (FAC) was used to supplement the culture medium for 24 hr prior to the acute RSL3 treatment defined above (i.e. 200 nM for 2 hr) (Fig. 1C i). Iron-loading alone dose-dependently increased lipid peroxidation reaching significance when the supplemented iron concentration was 50 µM or above (Fig. 1C ii) but did not compromise cell viability (Fig. 1C iii). In contrast, subsequent treatment with RSL3 only induced a modest further increase in lipid peroxidation (>50 µM FAC) (Fig. 1C ii) but a more dramatic change in cell death at concentrations > 50 µM FAC (Fig. 1C iii). Therefore, combining a pretreatment with 50 µM FAC for 24 hr followed by an acute RSL3 treatment for 2 hr had a synergistic effect in inducing lipid peroxidation without extensively compromising HMC3 cell survival.

To validate the ferroptotic nature of cell death in this model, we used Ferrostatin-1 (Fer-1) (10 µM) as a selective ferroptosis inhibitor and confirmed that it abolished lipid peroxidation and subsequent cell death (Fig. 1D). Finally, to exclude confounding cell death mechanisms such as apoptosis, we observed no cleavage of Caspase-3 (a crucial executioner enzyme in apoptotic cell death) except with the positive control, menadione (Fig. 1E).

Together, these data demonstrate that preloading of iron to HMC3 microglia primes ferroptotic death initiated through GPX4 inhibition, thereby providing a robust platform in which to further investigate initial molecular events preceding microglial ferroptosis.

### Iron-driven ferroptosis induces reactive oxygen species and endoplasmic reticulum morphological alterations in HMC3 microglia

Along with the well reported increased levels of Reactive Oxygen Species (ROS)^41^, there are also characteristic morphological changes to organelles^42^, particularly mitochondria^6,7^ and the endoplasmic reticulum^43^, associated with ferroptosis. To determine whether our HMC3 model exhibits such features, we initially used CellROX to image total ROS levels (Fig. 2A). Iron-laden cells displayed a modest elevation in total ROS compared to the DMSO control, while RSL3 alone or in combination with FAC increased ROS levels significantly (Fig. 2A i-ii). Importantly, this effect was reversed by co-treatment with Fer-1 and thus confirmed the ferroptotic nature of our model. We next used the MitoSOX dye on a flow cytometer to differentiate between levels of superoxide (Fig. 2B i) and total levels of ROS (Fig. 2B ii) generated by the mitochondria. HMC3 loaded with iron alone exhibited an increased level of mitochondrial superoxide but not of total ROS, suggesting that iron and RSL3 treatments modulate separate pools of ROS.

**Figure 2:**
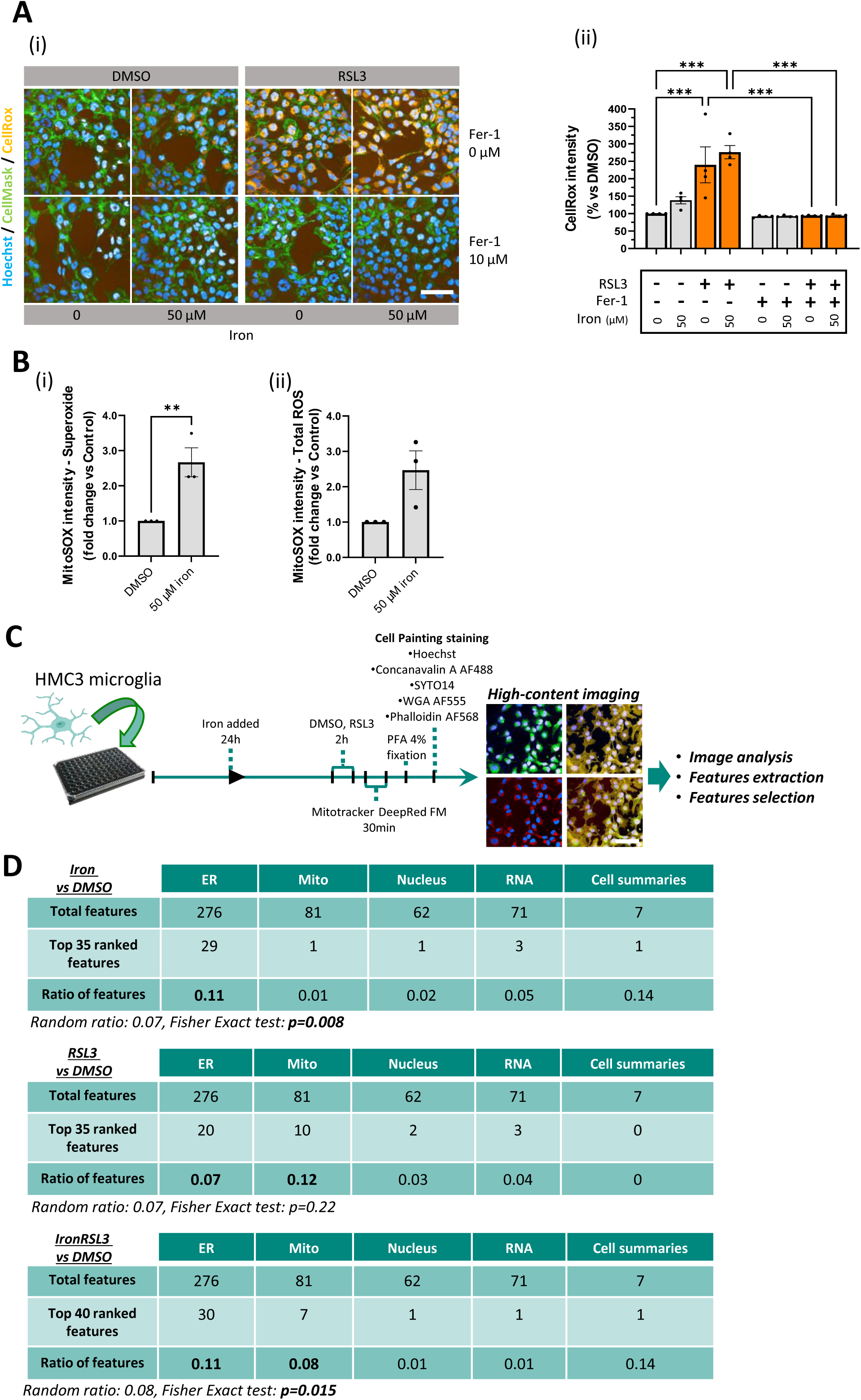
Iron-mediated cellular changes in HMC3. A) (i) Representative images of HMC3 cells stained with CellMask Green and CellRox Orange. The staining was performed after 24 hr iron-loading of the cells (50 µM FAC) followed by treatment for 2 hr with DMSO or RSL3 (200 nM) +/- Fer-1 (10 µM). Scale bar=100µm. (ii) Quantification of CellRox intensity for DMSO (grey) or RSL3 (orange). N=3, one-way ANOVA with Šidák’s test, ***p < 0.001. B) MitoSOX labelling and quantification using flow cytometry to detect (i) mitochondrial superoxide and (ii) total ROS. N=4, one-way ANOVA with Šidák’s test, **p < 0.00. C) Workflow of cell culture for the model and cell painting analysis. Scale bar=100µm. D) Contingency tables listing the numbers of most important features per organelle/compartment in the top 35-40 features, ranked by Random Forest classifier, to differentiate treatment conditions. Cell summaries were excluded from Fisher Exact Test as too few of these features were present.

To further characterise the impact of iron and RSL3 on organelle morphology, comprehensive phenotypic profiling was assessed using Cell Painting^44^ (Fig. 2C). The morphological features were initially extracted from Cell Painting imaging (Signals Image Artist) including all Intensity features, STAR morphology features and SER Texture features for the different organelles and then ranked using a Random Forest classifier to distinguish each condition from DMSO treated controls. Using the ratios of features compared to the Random Ratio (see Methods for calculations and Table 1 for top ranked features) for each organelle in the different treatment conditions, we identified that the ratio of ER features consistently exceeded the Random Ratio across all treatment conditions and thus was most likely attributable to iron-driven modifications. In contrast, the ratio of mitochondrial features only exceeded the Random Ratio in conditions where RSL3 was present. The Fisher Exact test determined that the distribution of organelles features differed from random chance in the iron-laden cells and in the iron-laden followed by RSL3 treatment but could not confirm non-random distribution in the condition with RSL3 only (Fig. 2D).

To explore possible mechanistic pathways underlying the ER morphological changes, key markers of the three Unfolded Protein Response (UPR) signalling branches (Sup Fig. 2A), levels of phosphorylated eIF2α (Sup Fig. 2B) and protein synthesis (Supp. Fig 2C) were all investigated. While direct changes to ER stress pathways were not observed (Sup Fig. 2A and B), protein synthesis rate was decreased to levels comparable with the positive control (CHX) in iron-laden HMC3 treated with RSL3, suggesting that shutdown of protein synthesis in cells succumbing to ferroptotic death could contribute to the ER morphology changes detected by Cell Painting profiling.

In summary, the identified phenotypic features in our human microglial HMC3 cell model confirm previously reported mitochondrial alterations only when cells are in full ferroptotic conditions^6^ but also reveal a prominent iron-driven modification to ER morphology that is independent of UPR signalling when cells are primed with iron. Together with the ROS evaluation, these results confirm that our model recapitulates the central molecular and cellular features of ferroptosis.

### Iron priming and RSL3 induction of ferroptosis increase sterol metabolic pathways

Ferroptosis is uniquely defined by the excessive accumulation of intracellular lipid peroxides. In past evaluation with non-CNS cell types this lipid peroxidation has predominantly been shown to originate from phosphatidylcholine (PC) and phosphatidylethanolamine (PE) phospholipids, however more recently other lipid forms including sterols have begun to be implicated as drivers of ferroptosis^45–47^. To unravel specific signatures of ferroptotic cell death associated with lipid peroxidation in our model, we performed multi-omics profiling of the transcriptome, proteome, and both the unoxidized and oxidized lipidomes (Fig. 3A). Before profiling each condition, samples were quality checked by confirming levels of lipid peroxidation and cell viability (Sup Fig. 3A) as well as key markers associated with ferroptosis: FTH, GPX4 and TfR1 (Sup Fig. 3B and C). In the RSL3 alone condition, we did not identify any significant changes in the lipid profile possibly to due to short time of RSL3 exposure. In iron-primed HMC3s, however, 121 upregulated and 58 downregulated differentially abundant lipids (DALs) were identified with Cer d18:2_22:0 being the most significant upregulated amongst several PC-, PE- and PS-PUFAs and their respective oxidised forms (Fig. 3B and Table 2). When cells were subsequently treated with RSL3, 199 upregulated and 26 downregulated DALs were identified with distinct accumulation of sterols including 7-oxo-cholesterol and 7α,27-dihydroxycholesterol. Ether phospholipids were also identified amongst DALs, 10 were upregulated and 3 downregulated in the iron-laden HMC3 while 17 were upregulated, including O-LPE, and 5 were downregulated in the iron-laden HMC3 subsequently treated with RSL3 (Fig. 3C and Table 2). While the ether phospholipids were not significantly affected by Fer-1 treatment, the sterol accumulation was partly reverted upon treatment with this ferroptosis inhibitor, with 12 downregulated DALs including cholesterol esters (Figs. 3B and C). A lipid class enrichment analysis confirmed that sterols were the second most enriched class after phosphatidylethanolamines upon iron and RSL3 treatment (Fig. 3D) and that oxysterols, cholesterols and cholesterol esters were in the top 5 restored lipid classes upon ferroptosis inhibition with Fer-1 (Fig. 3E i). Moreover, the relative abundance of specific lipid species decreased upon Fer-1 treatment was mainly PUFA containing phospholipids characterised by two or more double bonds in their fatty acyl chains (fatty acyl chains are indicated by the format C:n where C is the number of carbons in the chain and n the number of double bonds) alongside 7-oxo-cholesterol (Fig. 3E ii).

**Figure 3:**
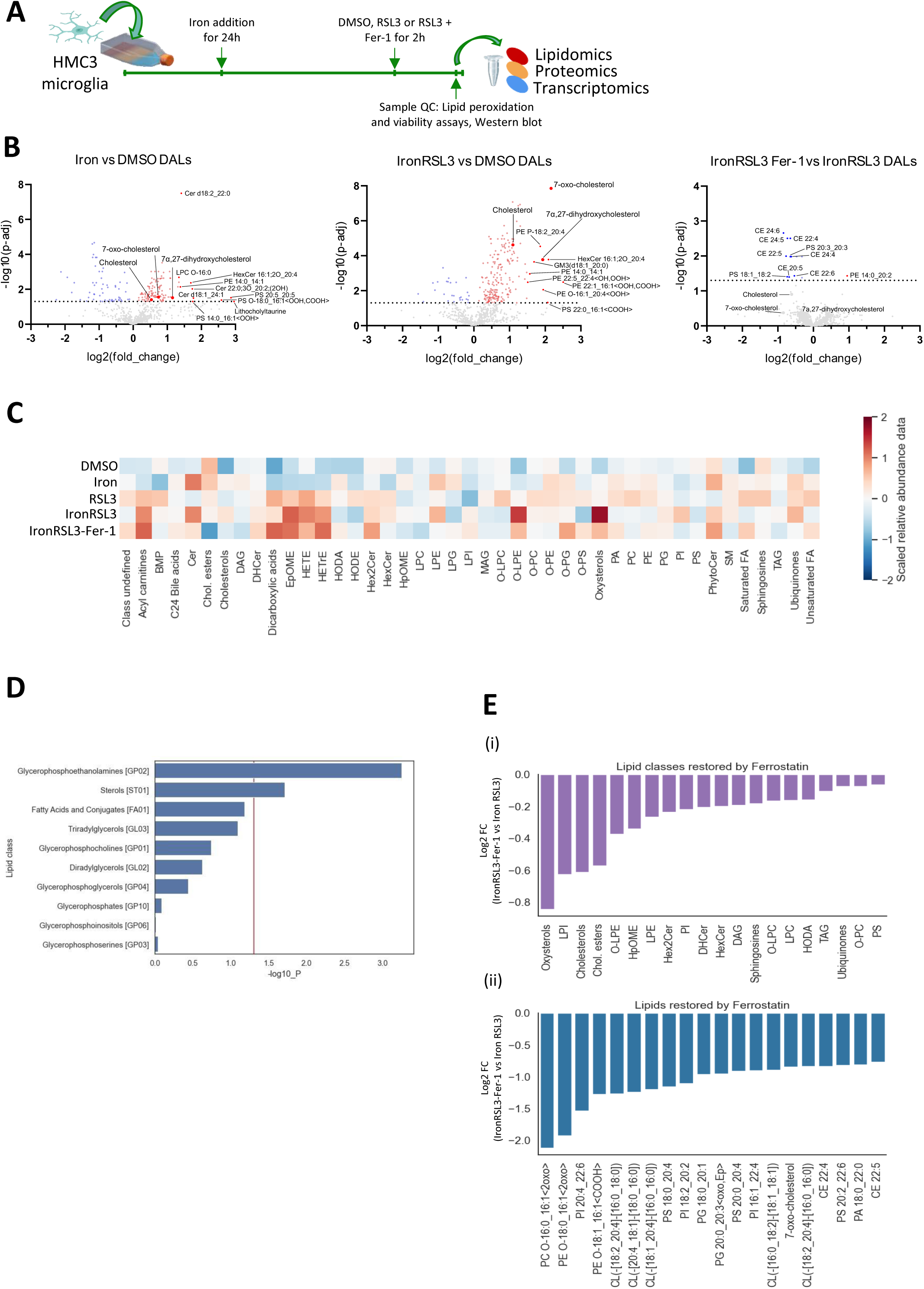
Lipidomic analysis of iron primed and RSL3 induced HMC3. A) Overview of sample collection process. B) Differentially abundant lipids represented on Volcano plots for all the treatments. The top 10 significant lipids, ranked by adj. p-value and log2 fold-change are highlighted alongside most significant sterols. C) Heatmap showing abundance of lipids averaged across replicates and LipidMaps classes. Lipid abundance data is on a unit-variance scale to make analytes comparable with each other. D) Lipid enrichment analysis barplot showing top 10 lipid classes upregulated in ferroptotic HMC3 (positive log2 fold change of IronRSL3 vs DMSO). Sub-classes obtained from LipidMaps. MWU test to identify differential lipids (p <0.05) and ORA performed on lipid classes). E) Barplots showing top 20 (i) LipidMaps classes and (ii) lipid species restored by Fer-1 treatment (negative log2 fold change of IronRSL3-Fer1 vs IronRSL3).

To identify whether profound changes of lipid classes and lipid oxidation would relate to dysregulation of regulatory pathways, the transcriptome and proteome were evaluated. Indeed, differentially expressed genes (DEGs) (Table 2) involved in the mevalonate pathway (e.g. HMGCS1, IDI1, SQLE, MSMO1 and DHCR7), cholesterol metabolism (e.g. ABCA7, LDLR, INSIG1, NPC2) and fatty acid metabolism (e.g. SCD and FADS2) were all significantly increased in HMC3 pre-loaded with iron and in HMC3 subsequently treated with RSL3 (Fig. 4A). Interestingly, HMGCS1, MSMO1, LDLR and INSIG1 were downregulated upon Fer-1 treatment suggesting that these may form part of a core gene signature of a pre-ferroptotic state in HMC3 (Fig. 4A). Spearman partial correlations further confirmed the involvement of cholesterol synthesis and uptake with oxidised lipid abundance in this ferroptotic model, whereby MSMO1 positively correlated with 7-oxo-cholesterol, HMGCS1 and LDLR positively correlated with the Arachidonic Acid derived oxylipin 11,12-DiHETrE, and INSIG1 correlated with an oxidised Ceramide Phosphoethanolamine (CerPE) (Fig. 4B). HMGCS1 and LDLR were also found upregulated among differentially abundant proteins (DAPs) in the proteome of HMC3 only pre-loaded with iron and in iron-laden HMC3 subsequently treated with RSL3 (Fig. 4C and Table 2), while HMGCS1 was downregulated after Fer-1 treatment (Fig. 4C). Finally, using both DEGs and DAPs for pathway analysis, we identified steroid and sterol metabolic processes as the top GO terms positively enriched in the HMC3 only pre-loaded with iron and in iron-laden HMC3 treated with RSL3. Transcriptionally, these GO terms were also the most negatively enriched upon Fer-1 treatment (Fig. 4D).

**Figure 4:**
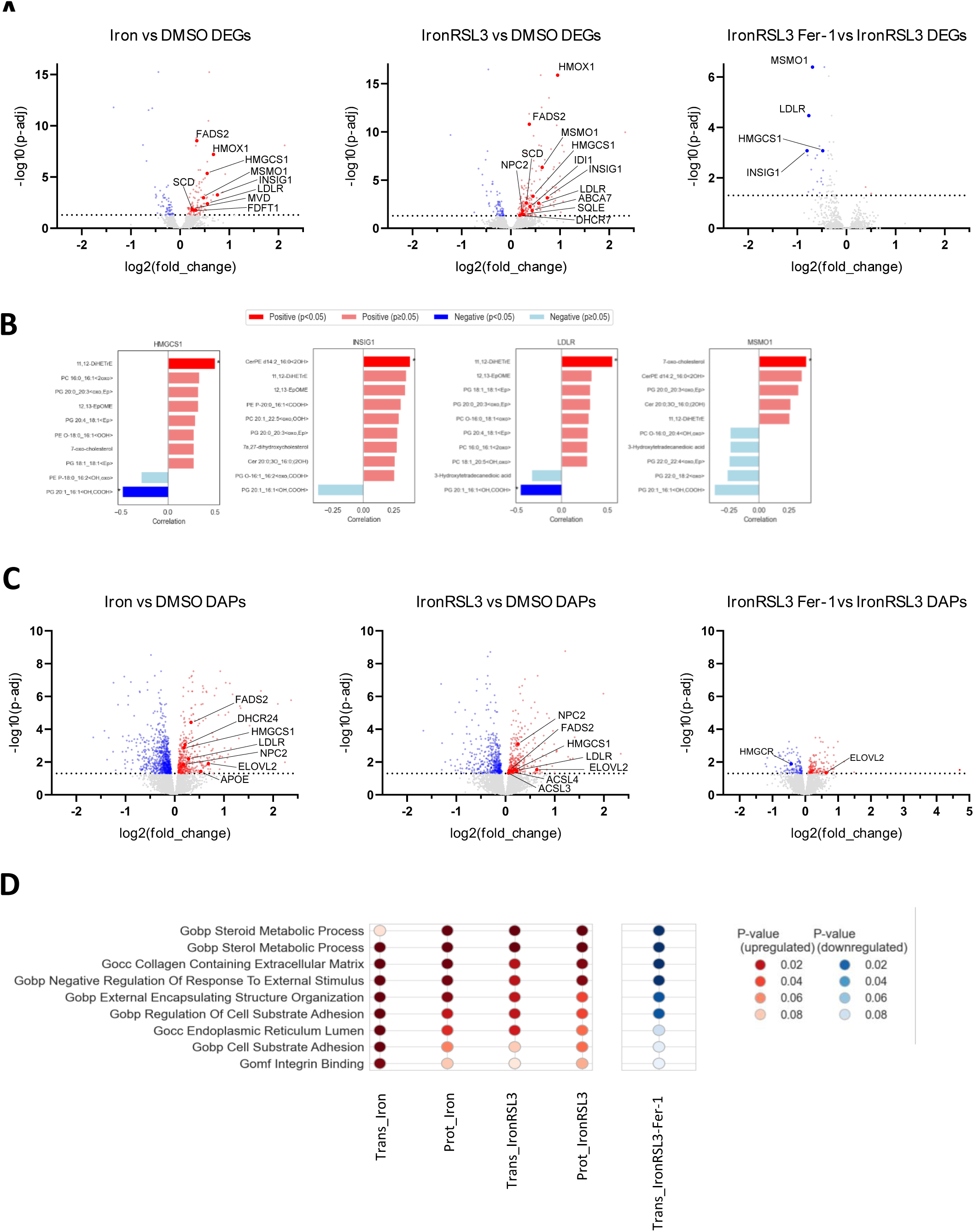
Transcriptomic and proteomic analysis of iron primed and RSL3 induced HMC3. A) Differentially expressed genes (DEGs) represented on Volcano plots for all the treatments. DEGs related to metabolism of fatty acids and cholesterol are highlighted. B) Spearman partial correlations between the DEGs involved in cholesterol metabolism and oxidized lipid abundance, with treatment as a covariate. Significant positive (red) or negative (blue) correlations are highlighted with an asterisk (*p < 0.05). C) Differentially abundant proteins (DAPs) represented on Volcano plots for all the treatments. DAPs related to fatty acids and cholesterol metabolism are highlighted. D) Common significant pathways between transcriptomics and proteomics datasets (Gene Ontology, KEGG, and Reactome, significance set at p ≤ 0.1). No significant pathways for the ferroptosis inhibition condition were found in the proteomic data.

In summary, evidence supports that in addition to modulation of PUFAs the HMC3 model in a pre-ferroptotic state also elevates metabolic pathways related to sterol with a subsequent increase in levels of oxysterols.

## Discussion

Priming of HMC3 microglia with iron (24 hr) before inhibition of the key ferroptotic regulator GPX4 (RSL3 treatment for 2 hr) provides a sub-lethal cellular model of ferroptosis that enables evaluation of time dependent changes. Key molecular features in microglia included increased lipid peroxidation, increased ROS levels and marked morphological alterations in the endoplasmic reticulum. Lipid signatures distinct from PUFAs were identified, with this pre-ferroptotic state accumulating sterols and oxysterols as well as a concomitant upregulation of genes and proteins involved in sterol metabolism^38,39^. Findings presented here also reveal a distinct gene signature already present in iron-laden HMC3, specifically HMGCS1, MSMO1, LDLR, and INSIG1, that coincides with an accumulation of sterol lipids. This was sustained in iron-laden HMC3 induced for ferroptosis by RSL3 together with a notable increase of oxysterols and ROS levels. The upregulation of genes involved in cholesterol biosynthesis and uptake suggests an early cellular adaptation aimed at modulating membrane composition to potentially protect against ferroptotic cell death. A similar mechanism has been reported in HEK293T cells whereby subsequent genetic or pharmacological inhibition of DHCR7, a critical regulator in the terminal steps of cholesterol synthesis encoding 7-dehydrocholesterol reductase, causes accumulation of 7-dehydrocholesterol and increased lipid peroxidation to sensitise the cell for ferroptosis^16^. While elevated cholesterol levels may serve as an intermediary protective mechanism by buffering cells against the damaging effects of rising ROS, this in turn creates a substrate pool that, upon oxidation, will promote cell death. In a similar vein, we observed a cumulative upregulation of the antioxidant *HMOX1*^48^ between the condition with iron alone and iron combined with RSL3, suggesting that microglia activate parallel defensive mechanisms against ferroptosis. Therefore, this model would be suited for further investigations to identify targets promoting resistance against ferroptosis at the node between antioxidant and lipid metabolism pathways.

Likewise, our modelling delineates a link between the lipid and ROS changes associated with iron loading of microglia and morphological changes to the ER that is non UPR-related. This underscores the critical roles that organelles such as the ER, mitochondria, and peroxisomes serve as regulatory hubs of lipid and ROS metabolism. While mitochondrial alterations have been strongly associated with ferroptosis^49^, growing evidence demonstrates that other organelles are specifically regulated as cells undergo ferroptosis^9,12,43^. This study’s findings confirm that these mitochondria morphological changes occur in microglia but only when cells are fully undergoing ferroptotic induction (i.e. upon RSL3 treatment), whereas ER morphological changes are prevalent even at a pre-ferroptotic stage when intracellular iron is accumulating. The dynamic interplay between these organelles in relation to lipid metabolism appears central to maintain cellular homeostasis and identifying novel modulatory signalling pathways could further enhance our understanding of early stages in cellular susceptibility to ferroptotic cell death.

Finally, the identification of a core gene signature associated with cholesterol metabolism comprising HMGCS1, MSMO1, LDLR, and INSIG1 in our model could pave the way for demonstrating a direct association between this pathway and the role of ferroptosis in diseases where microglia are heavily implicated in neurodegeneration. MSMO1 alone emerges as a potential biomarker in PD^50^. The involvement of ferroptosis in PD is increasingly recognised, particularly through an interplay between ether-phospholipids and alpha-synuclein^47^. The familial PD A53T mutation in alpha-synuclein has also recently been reported to increase sensitivity to ferroptotic death, including in microglia^51^. Single nuclei RNA-sequencing of microglia subpopulations from sporadic PD brains has identified five subpopulations amongst which two display upregulated DEGs involved in the UPR and oxidative stress pathways^52^. One of these two microglial subpopulations of interest also harboured upregulated DEGs associated with immune response and iron storage genes (i.e. FTL and FTH1). These different subpopulations could thus represent microglia at different ferroptotic stages in sporadic PD. While primary and iPSC-derived microglia would offer enhanced physiological relevance to explore potential ferroptotic associations in these subpopulations of microglia, their high variability and complex culture requirements pose significant challenges for longitudinally evaluating microglial ferroptosis. In contrast, the HMC3 immortalised microglial cell line provides genetic consistency and ease of maintenance as well as a capability to robustly model disease relevant mechanisms^53^. Overall, it is proposed that these microglial subpopulations found in sporadic PD could align with the different ferroptotic stages modelled in HMC3. Combining our robust microglial model of ferroptosis with high throughput screening methods such as CRISPR or compound libraries, as well as multiomic approaches, should pave the way to identifying the early molecular events that underpin microglial vulnerability to ferroptosis, including dissecting the interplay between lipid composition of membranes and associated genes. These could then be validated and further refined with iPSC-derived patient cells to investigate how microglia may contribute to neurodegeneration in PD and other neurodegenerative diseases.

## Materials and Methods

### Chemical treatments and supplementation

Erastin (MedChemExpress, HY-15763), RSL3 (Sigma, SML2234), Ferrostatin-1 (Fer-1; Sigma-Aldrich, SML0583), and Ferric Ammonium Citrate (FAC; Sigma-Aldrich, F5879) were used to model ferroptosis.

### Cell culture

Human microglia HMC3 cell line was purchased from ATCC (CRL-3304) and cultured in MEM containing Earl’s Salts and glutamine (Gibco, 31095-029) supplemented with 10% FBS (Gibco, 16140-063), 1% penicillin and streptomycin (Gibco, 15140-122) at 37°C and in a humidified atmosphere containing 5% CO2. Cells were seeded at a density of 1.56 x 10^4^/cm^2^ for all assays and when required for sample collection or subculturing, they were dissociated with TrypLE (Gibco, 12604013).

### Lipid peroxidation and cell viability

HMC3 cells were dissociated with TrypLE and transferred to a V-bottom 96-well plate for incubation with BODIPY 581/591 C11 (10 µM, Invitrogen, C10445) at 37°C for 30 min. DRAQ7 (3 µM, Invitrogen, D15106) was then added to cells before loading the plate into the Attune NxT Flow Cytometer (ThermoFisher Scientific). Lipid peroxidation levels were determined by measuring the percentage of HMC3 positive for the oxidised form of BODIPY 581/591 C11 among the total HMC3 population. Viability was determined by measuring the percentage of the DRAQ7 positive population among total cell population. Controls without dye were used to set the mean intensity fluorescence threshold for both dyes.

### Cleaved Caspase 3 immunocytochemistry

HMC3 cells following treatment with Iron and RSL3 as previously described, or with 100 µM Menadione (MilliporeSigma, M5625) were fixed in 4% PFA for 15 min, permeabilised and blocked with 5% BSA in PBS + 0.1% Triton X-100 for 30 min. Cells were then incubated overnight at 4°C with Cleaved Caspase-3 (Asp175) antibody (Cell Signalling, 9661) in Dilution Buffer (1% BSA and 0.1% Triton X-100 in PBS, 1/400). After 3x washes with PBS, the cells were incubated for 1 h at room temperature with the corresponding Alexa-conjugated secondary antibodies (1/1,000 in Dilution Buffer) and counterstained with Hoechst (1/10,000). Immunofluorescence images were acquired on the Opera Phenix High-Content Imaging System (Revvity) and images were analysed using Signals Image Artist.

### ROS measurement

Total ROS levels were measured by staining cells for 30 min with CellROX Orange (ThermoFisher Scientific, C10443, 1/500) and CellMask Green (ThermoFisher Scientific, C37608, 1/1,000) and counterstained with Hoechst (1/10,000). Cells were imaged on an Opera Phenix High-Content Imaging System (Revvity) and quantification was performed with Signals Image Artist. MitoSOX Red (ThermoFisher Scientific, M36008, 500 nM) was also used to measure total ROS and mitochondrial superoxide. The cells were stained for 30 min with MitoSOX and fluorescence was measured on the Attune NxT Flow Cytometer. Quantification of superoxide and general ROS was based on the respective fluorescence peaks, excitation at 396 nm and fluorescence detection by VL3 filter for superoxide and excitation at 500 nm with fluorescence detection by BL2 filter for the total ROS.

### Viability time-lapse recording

HMC3 were seeded in a 6-well plate (Corning, 3516) and treated with DMSO or RSL3 (200 nM) for 2 hr. After wash out with PBS, fresh medium was added and the cells were placed in the Incucyte S3 Live-Cell Analysis Instrument (Sartorius) for time-lapse recording, at 37°C and in a humidified atmosphere containing 5% CO2. Phase-contrast images were acquired every 45 min for a total duration of 24 hr. Cellular confluence was used as primary metric for survival.

### Cell Painting

HMC3 were seeded in a PDL-coated PhenoPlate 96-well microplate (Revvity, 6055500). Following respective treatment, Mitotracker Deep Red FM was added into each well and incubated for 30 min at 37°C. Cells were then fixed in 4% paraformaldehyde (PFA) for 15 min and permeabilized with 0.1% Triton-X100 in PBS for 15 min. Hoechst 34580 (5 µg/mL), Concanavalin A-AF488 (50 µg/mL), Wheat Germ Agglutinin-AF555 (3 µg/mL), AF568-Phalloidin (6.5 µM) and SYTO 14 (300 ng/mL) from the Image-iT™ Cell Painting Kit (ThermoFisher Scientific, I65000) were diluted in HBSS with 1% BSA and applied to the cells for 30 min before wash out and sealing of the plate. Cells were imaged on an Opera Phenix High-Content Imaging System (Revvity, 20 ROI per well, 40X, 3 Z-planes). Analysis was performed with Signals Image Artist. A broad set of quantitative features describing morphology (STAR), texture (SER), and fluorescence intensity was extracted for the different organelles. After removing highly collinear features to reduce redundancy, a total of 497 features was identified for downstream analysis. Among these, the top predictive features distinguishing ferroptotic HMC3 from healthy control cells were ranked by Random Forest classifier and significance was tested with a Fisher Exact test to confirm non-random distribution. The Random Ratio was calculated by dividing the number of top ranked features selected (i.e. 35 for the Iron and the RSL3 conditions and 40 for the IronRSL3 condition) over the total number of analysed features (497). For each treatment condition, the ratio of features for each organelle was calculated by dividing the number of features associated with the corresponding organelle amongst the top ranked ones by the total number of features analysed for this organelle (e.g. the ratio of ER features in Iron vs DMSO corresponds to 29 features amongst the top ranked ones divided by 276 total ER features).

### Quantitative RT-PCR

TaqMan Gene Expression Assays (Applied Biosystems, 4331182) and TaqMan Fast Advanced Mastermix (Applied Biosystems, 4444558) were used on a QuantStudio 7 Flex system (ThermoFisher Scientific). The following TaqMan assays were used for human gene expression analysis: ATF4 (Hs00909569_g1), ATF6 (Hs00232586_m1), XBP1 (Hs00231936_m1), GPX4 (Hs00989766_g1). Relative gene expression was determined via the Comparative CT method normalised to at least 3 housekeeping genes (ACTB (Hs01060665_g1), B2M (Hs00187842_m1), GAPDH (HS02786624_g1), TBP (Hs00427620_m1)).

### Western Blot

Following respective treatment, HMC3 cell pellets were collected from 6-well plates (Corning, 3516) and resuspended in RIPA lysis buffer (Fisher, 89900) with added protease and phosphatase inhibitors (Sigma-Aldrich, 4693116001 and 4906837001 respectively) and incubated on ice for 15 min with regular vortexing. The samples were then centrifuged at 14,000 *g* for 15 min at 4°C and the supernatant collected. Protein concentration was measured with the Pierce^TM^ BCA Protein Assay Kit (Fisher, 23227). Samples were prepared to a final concentration of 1 µg/µL with 1X NuPAGE LDS sample buffer (Invitrogen, NP0007), 1X NuPAGE reducing agent (Invitrogen, NP0009) and deionised water. Samples were heated at 70°C for 10 min. Samples were loaded into NuPAGE 4-12% Bis-Tris 10-well gel (Invitrogen, NP0335BOX) alongside Chameleon Duo protein ladder (Li-COR, 928-6000). Gels were run in 1X MES running buffer (Invitrogen, NP0002) at 150V constant for 80 min and transferred to a nitrocellulose membrane using the Bio-Rad Mini Trans-Blot in 1L of chilled, pre-prepared transfer buffer (1X Bio-Rad Tris/Glycine Transfer Buffer (BioRad, 1610771) with 10% Methanol), at 100V/0.35A constant for 1 hr. Membranes were washed in PBS and blocked using PBS Intercept Blocking Buffer (LICORbio, 927-70001) for 1 hr at room temperature with rolling. Primary antibody solution diluted in blocking buffer + 0.2% Tween 20 was applied to the membranes incubated at 4°C overnight with rolling: GAPDH anti-mouse (Abcam ab8245, 1/1,000) GPX4 anti-goat (Invitrogen 11336523, 1/500), FTH anti-rabbit (Abcam ab65080, 1/500) and TfR1 anti-mouse (Invitrogen 13-6800, 1/1,000). After incubation, membranes washed 3 times with 0.1% Tween 20 in PBS before adding respective secondary antibodies for 1 hr at room temperature in the dark (IRDye 800CW donkey anti-goat (LI-COR 926-32214), IRDye 800CW donkey anti-rabbit (LI-COR 926-32213) and IRDye 800CW donkey anti-mouse (LI-COR 926-32212), all with a dilution of 1/20,000). Before imaging, membranes were washed in 0.1% Tween 20 three times and one time in PBS, and acquisition was performed on a LI-COR Odyssey and analysed with Image Studio Lite software (Version 6.0.0.28).

### SUnSET assay

HMC3 were seeded in a non-PDL-coated PhenoPlate 96-well microplate (Revvity, 6055302). Cells were treated for 2 hr as previously described, with Cycloheximide treatment (CHX, 50 µg/mL) included as positive control for inhibition of the protein synthesis, before addition of Click-iT OPP reagent (ThermoFisher, C10458, 20 µM) for 30 min. Cells were then fixed (PFA 4%), permeabilised (0.1% Triton X-100) and incubated 30 min at room temperature with Click-iT OPP Reaction Cocktail (prepared according to Click-iT OPP manufacturer’s instructions). Subsequent steps were all performed at room temperature and protected from light. Wells were washed with 50 µL of Click-iT Reaction Rinse Buffer before adding 100 µL per well of HCS Nuclear Mask Blue Stain for 30 min. All wells were washed with PBS before sealing the plate and imaging on an Opera Phenix High-Content Imaging System (Revvity) to capture the Click-iT OPP signal (Alexa Fluor 647) corresponding to newly synthesized proteins.

### HTRF assay

HMC3 cells were cultured in 96-well culture plates and treated as described previously, including Thapsigargin treatment (1 µM) as positive control. After treatment HTRF Human and Mouse Phospho-eIF2α (Ser52) kit (Revvity, 64EF2PEG) and Total eIF2α kit (Revvity, 64NEFPEG) were used. Following manufacturer’s recommendations, cells were lysed with 50 µL per well of lysis buffer for 30 min at room temperature with shaking. 16 µL of each lysate were transferred to two separate white detection plates (Corning, 3610) with addition of 4 µL of the corresponding premixed antibody solution. Plates were sealed and incubated at room temperature overnight before reading the fluorescence at 665 nm and 620 nm on a PHERAstar microplate reader (BMG LABTECH). The ratio of signal at these two wavelengths was used to measure the proportion of phosphorylated eIF2α.

### Collection of samples and QC for OMICs

Samples for OMICs were treated as previously described for induction/inhibition of ferroptosis. Cells were dissociated using TrypLE (Gibco), counted and pelleted with a small fraction spared for QC analysis. Pellets were washed in PBS containing 5 mM PMSF and 5 mM BHT to prevent protein degradation and oxidation. Cells were divided equally into three tubes for each sample for separate OMICs analysis in LoBind tubes (Eppendorf, 0030108116), pelleted and flash frozen in dry ice. QC for each batch of samples included BODIPY/DRAQ7 assay for lipid peroxidation, cell viability and western blotting for protein expression of GPX4, TFRC and FTH1.

### Bulk RNA-sequencing and Transcriptome analysis

Frozen cell pellet samples (5 conditions, 3 biological replicates) were submitted to Genewiz (Azenta Life Sciences) for bulk RNA sequencing. Total RNA quality was assessed, and messenger RNA was enriched using poly(A) selection prior to library construction. Libraries were sequenced on an Illumina® NovaSeq™ platform with a 2 × 150 bp pairedLJend configuration to an average of ∼30 million reads per sample. Genewiz guaranteed ≥85% of bases at Q30 or higher and supplied FASTQ files and a QC report. All samples passed QC. Reads were trimmed, filtered, and mapped to the reference genome by Genewiz using a standard preprocessing pipeline. Gene counts obtained from FASTQ files for all the genes were then analysed for detection of DEGs between DMSO condition and the different ferroptotic conditions using R limma, with experimental batch as a covariate. Genes with FDR-adjusted p-value ≤ 0.05 were considered DEGs.

### Proteomics

#### Protein Extraction, Quantification and Digestion

Cell lysis was achieved using the commercial LYSE buffer supplied in the In-StageTip digestion (iST) sample preparation kit (PreOmics). Lysis was promoted by sonication in a water bath sonicator for 15 min at 4^∘^C. Following cell lysis, protein concentration was determined using the Pierce^TM^ BCA Protein Assay Kit - Reducing Agent Compatible (ThermoFisher Scientific) according to the manufacturer’s instructions. 50 μg of proteins from each sample were used for subsequent proteomics digestion using the iST kit (PreOmics) following the manufacturer’s protocol. Briefly, 50 μg of protein were loaded onto the StageTips, and proteins were denatured by heating the tips at 95°C for 10 min with agitation at 1,200 rpm. Supplied trypsin/LysC DIGEST mixture was subsequently added, and samples were incubated at 37°C for 3 hr with shaking at 1,200 rpm. Digestion was halted by adding the supplied STOP buffer, and the resulting peptide mixture was cleaned by passing the supernatant through the provided filter cartridge. Peptides were eluted twice with the ELUTE buffer, vacuum-dried, and reconstituted in LC-LOAD buffer. Finally, 400 ng of peptide from each sample was loaded onto an Evotip Pure tip (Evosep) according to the manufacturer’s specifications and stored in 0.1% formic acid until mass spectrometry (MS) analysis.

#### Liquid Chromatography (LC)−MS/MS Analysis

LC-MS analysis was performed on an Evosep ONE LC system (Evosep Biosystems) connected to a timsTOF fleX MALDI-2 mass spectrometer (Bruker). The Evosep ONE was operated in the Whisper Zoom configuration, and separation was achieved using the Whisper Zoom 40 SPD method. Elution occurred over an Aurora Elite column (15 cm×75 μm×1.7 μm, IonOpticks), which was maintained at 50°C. The mobile phase consisted of Buffer A (0.1% formic acid in LC-MS grade water) and Buffer B (0.1% formic acid in acetonitrile). Eluted peptides were detected in dia-PASEF mode using the timsTOF fleX MALDI-2 mass spectrometer (Bruker) operated in positive ion mode equipped with a captive spray ion source. The capillary voltage was set at 1500 V with a capillary temperature of 180°C. The MS1 full scan ranged from m/z 100 to m/z 1700. dia-PASEF methods were configured for a 1.80 s total cycle time, utilizing 16 MS/MS PASEF cycles with fixed 26.0 Da isolation windows spanning the mass range from 400 m/z to 1200 m/z. Ion mobility separation was conducted over a 1/K0 range between 0.60 V⋅s/cm2 and 1.60 V⋅s/cm2, with a 100.0 ms accumulation and ramping time. A 1.0 Da mass overlap and 32 mass steps per cycle were applied. Collision energy was dynamically ramped across the ion mobility range to ensure comprehensive peptide fragmentation.

The MS RAW-files were analysed using Spectronaut v19.7.250203.62635 (Biognosys) with directDIA+ using default search settings. The database search was performed against a FASTA file containing the human proteome (SwissProt, 20,406 sequences, downloaded January 2025). Carbamidomethylation was set as a fixed modification, and N-terminal acetylation and oxidation of methionine were set as variable modifications. Trypsin/P was specified as the cleavage enzyme with a maximum of two missed cleavages.

The mass tolerance was set to dynamic in both MS1 and MS2 levels with a correction factor of 1. A Q value of 1% against mutated decoys was applied to filter identifications with a false-discovery rate (FDR) level of 0.01 at both the peptide and protein level. Quantification was performed using the automatic setting and normalized with the integrated cross-run normalization feature unless otherwise specified.

Proteomics data exported from Spectronaut was filtered such that triplicates with one missing value out of three measurements were imputed with a value drawn from the normal distribution that was based on the other two measurements (R scImpute^54^). Triplicates with two or three missing values were designated as missing not at random and were imputed with a value drawn from a normal distribution with mean down-shifted by 1.6 x SD (R tImpute^54^). Differential proteins were computed using R limma (R version 4.4.1) with eBayes moderation, and with triplicates handled by setting sample ID, which was common to each triplicate, as blocking factor. Correction for testing of multiple hypothesis was by the Benjamini & Yakutieli method with alpha = 0.05^55^ The thresholds for significantly different proteins were log2 fold-change < 0.5 and adjusted p-value ≤ 0.05, were considered as differentially abundant proteins (DAPs).

### Lipidomics

The harvested cells were resuspended in 100 µL of LC-MS grade water (Optima^TM^ Water, Fisher Scientific) and subjected to extraction of lipids by two-phase MTBE-Methanol liquid extraction protocol^56^. Internal standard mixture composed of SPLASH Lipidomix I (Avanti Research), 17:0 lyso PS-d5 (Avanti research, Cat No. 858148), 17:0 lyso PG-d5 (Avanti Research, Cat No. 858129), 19:0 lyso PI-d5 (Avanti Research, Cat No. 850108), including oxidized lipids standards PAzePC (Avanti Research, Cat. No. 870600), POVPC (Avanti Research, Cat No. 870606), and 5-oxoETE-d7 (Cayman Chemical, 334250) was added to each sample for quantification. Briefly, 200 µL of methanol (Optima^TM^, Fisher Scientific) was added to each sample and subjected to sonication in ice-bath (15 min) for cell lysis. Then, 660 µL of MTBE was added for non-polar phase extraction, and samples were incubated at room temperature for 30 min on a shaker. Phase separation was induced using 165 µL LC-MS grade water. Samples were centrifuged at 13,000*g* for 10 min at 4°C. Upper phase comprising non-polar lipids were collected in new microfuge tubes and dried under a stream of nitrogen gas. Dried lipid-containing fractions were resuspended in 100 µL of diluent 60:35:5 (Acetonitrile:2-propanol:Water) and subjected to LC-MS data acquisition. A pooled extract from each sample was used as technical quality control and analysed at regular intervals during the sample acquisition. All extraction solvents were supplemented with 100 µM BHT (Butylated Hydroxy Toluene, Sigma W218405).

Reconstituted lipid samples (3 µL) were injected using a chilled UPLC autosampler (Waters Corporation, Milford USA) and subjected to reversed-phase chromatography through ACQUITY CSH C18 column (2.1 x 100 mm, 1.7 µM, Waters, Milford, USA). Samples were eluted through a gradient of increasing solvent B (90:9:1 2-Propanol:Acetonitrile:water with 0.1% formic acid, 10 mM ammonium formate, and 100 µM BHT) from 15% to 99% over 12 min, pumped by a binary solvent manager (Waters, Milford, USA). The column was re-equilibrated for 3 min with solvent A (60:40 Acetonitrile: water with 0.1% formic acid, 10 mM ammonium formate, and 100 µM BHT). The eluted lipids were detected using a coupled high-resolution mass spectrometer SCIEX 6600 triple Tof (AB Sciex, Foster City, USA) operated using Analyst TF (v 1.8.1) software. Data was acquired in both positive and negative modes using an information-dependent acquisition with source parameters (400°C temperature and ± 4500 V) collected over 100-1500 m/z mass range.

Acquired LC-MS/MS lipidomics data was subjected to peak identification and annotation using Progenesis QI software (Non-linear dynamics, USA). The peaks were matched to an internally curated lipid database using search parameters described previously^57^. Each m/z peak was annotated to a lipid species based on MS/MS fragmentation, ± 10 ppm mass error, and retention time validation. Peaks with high signal-to-noise ratio and reproducibility across 3 biological replicates were only considered. Annotated lipid data output was then processed and normalized using R and Microsoft excel. Annotated lipidomics data was normalized to total ion abundance and lipid class-specific changes were normalized using specific deuterated internal standard. All values were also normalized to sample input using respective protein amounts. A Limma model was applied to the processed lipidomics abundance data, with experimental batch as a covariate. Lipids with fold change ±1.5 and FDR-adjusted p-value ≤ 0.05 were considered differentially abundant lipids (DALs).

Lipid abundance data was unit-variance scaled prior to heatmap visualisation. Lipid abundances were averaged across samples and LipidMaps lipid classes, resulting in a class-level heatmap.

### Pathway analysis

Pathway analysis was performed using the over-representation analysis method, implemented using the GSEApy Python package (v1.1.9). The background set contained all genes or proteins profiled in the experiment. Human Gene Ontology, KEGG, and Reactome databases were jointly used to compute pathway enrichment. Input differential metabolites or proteins were split into up and down-regulated sets based on the sign of their fold change, and pathway enrichment was computed separately for up and down-regulated analytes. Pathways with FDR-adjusted p-value ≤ 0.05 were considered significant.

### Correlation analysis

Spearman Partial correlations between the genes of interest and oxidised lipids were computed using the pingouin Python package (v0.5.5) partial_corr function. Treatment group was added as a covariate when computing partial correlations, and significant correlations were considered those with Spearman p-value ≤ 0.05.

### Statistical analyses

Results were expressed as the mean ± S.E.M. Sample size for each experiment is stated in the figure captions. Statistical analyses were performed using one-way or two-way ANOVA and post-hoc tests for multiple comparisons. Minimum statistically significant differences were established at p-value < 0.05. Non-statistically significant differences are not shown in the graphs.

---

**Supplementary Fig. 1:**
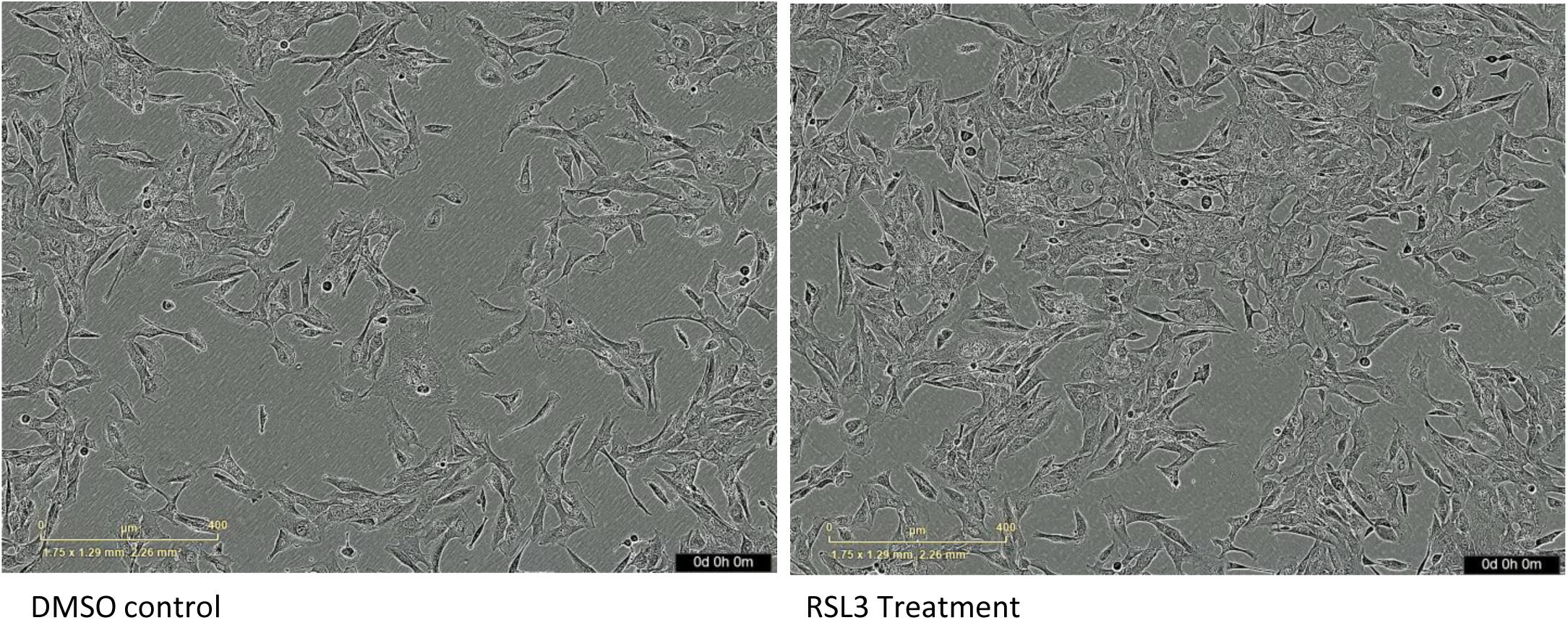
Cell viability over 24 h after acute RSL3 treatment. Incucyte recording of HMC3 cells post treatment. The cells were treated with DMSO or RSL3 (200 nM) for 2 hr, washed out and replenished with fresh medium for the rest of the recording. Frames were taken every 45 min for 24 hr.

**Supplementary Fig. 2:**
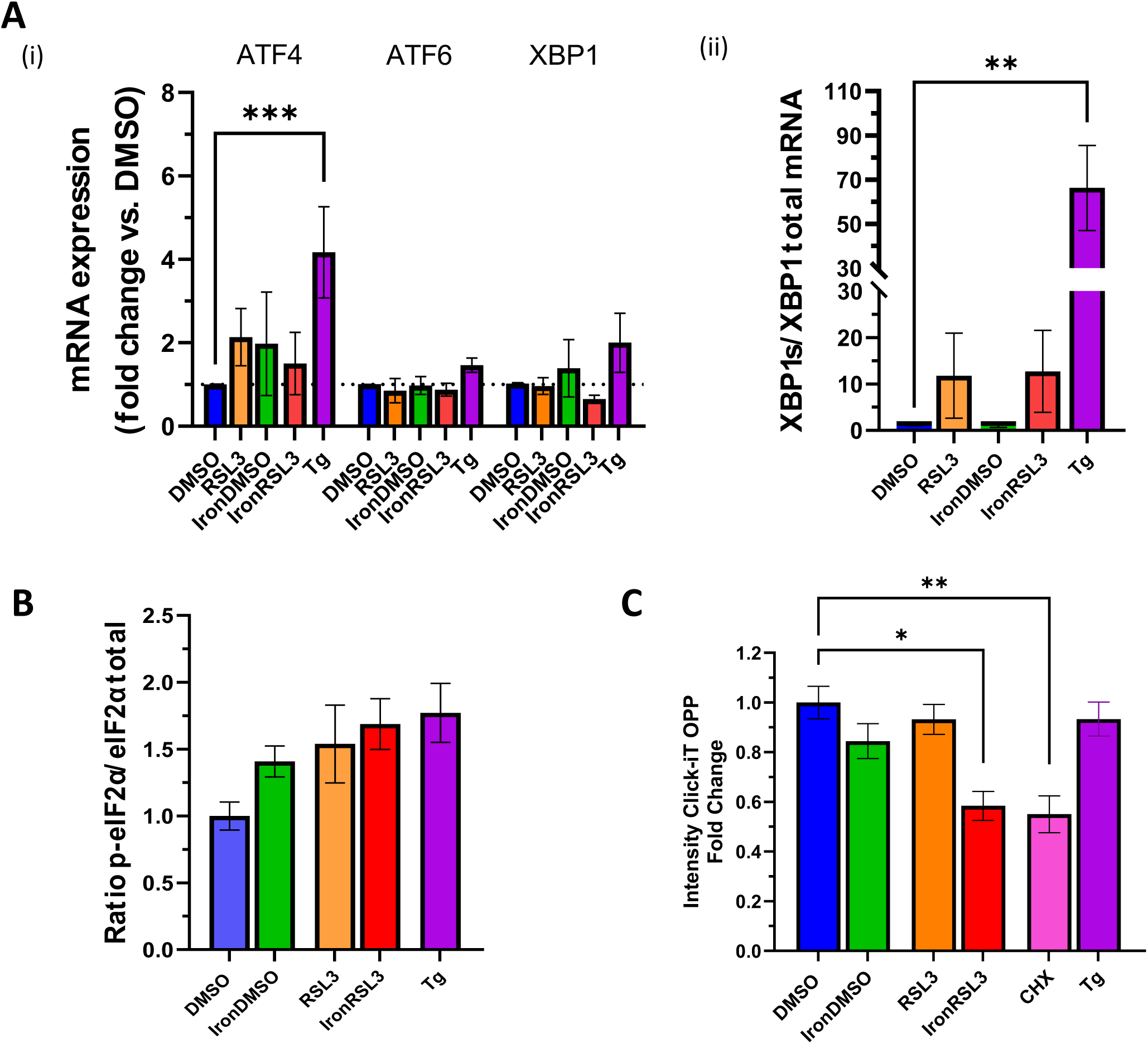
ER stress assessment. A) Measurement of (i) the mRNA expression of key markers of the UPR by qPCR (ATF4, ATF6 and XBP1) and of (ii) the ratio of spliced XBP1 over its unspliced form by qPCR following treatment, versus DMSO-treated controls. N=3, one-way ANOVA with Šidák’s test, **p < 0.01, ***p < 0.001. B) Quantification of phosphorylated eIF2α measure using HTRF method after treatments. N = 3, one-way ANOVA. C) Measurement of the rate protein synthesis in HMC3 after treatments. N=3, one-way ANOVA with Šidák’s test, *p < 0.05, **p < 0.01.

**Supplementary Fig. 3:**
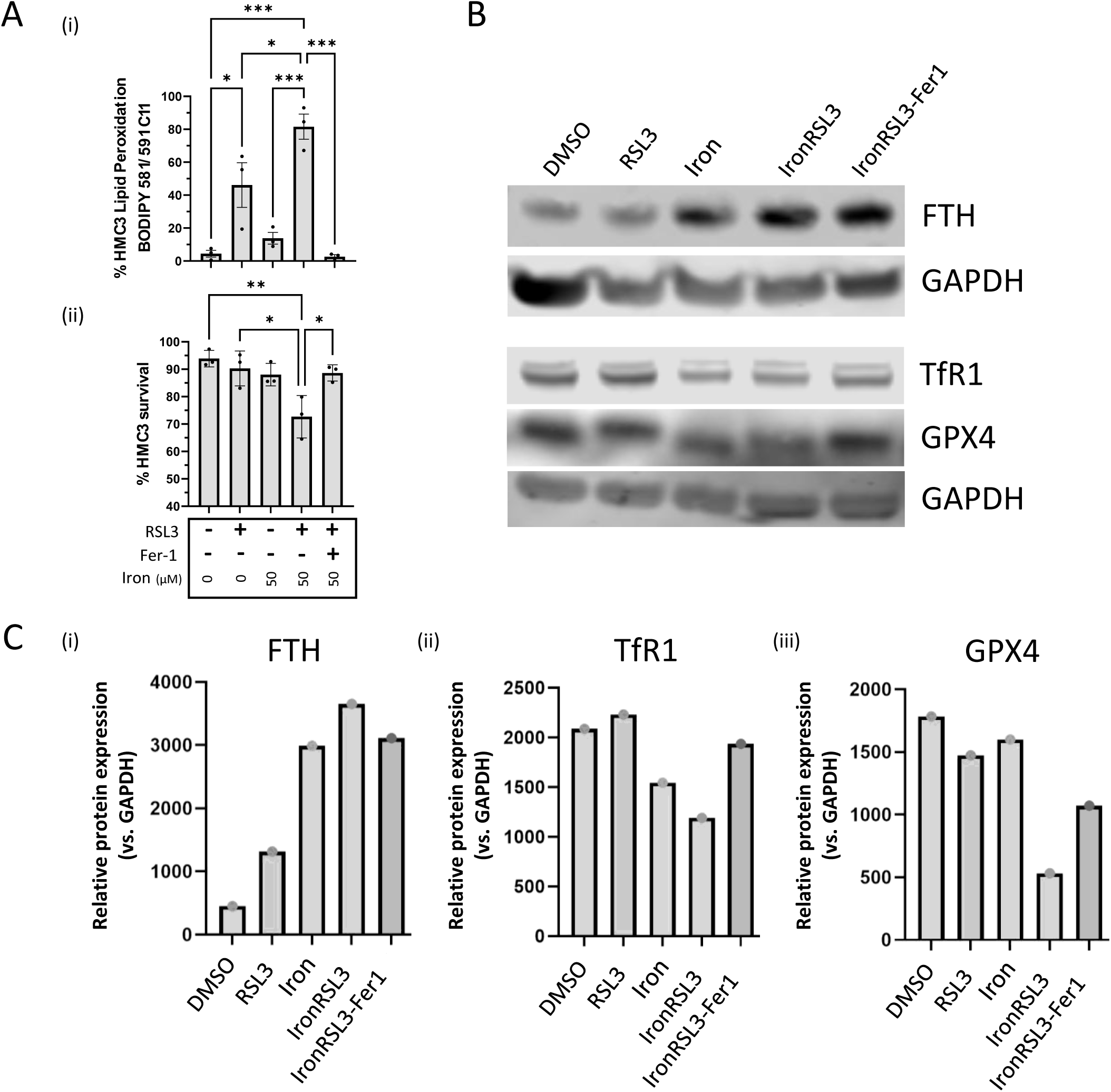
Quality control for multiome profiling. A) (i) Cytofluorimetric assessment of lipid peroxidation and (ii) cell viability. Three different batches including five treatment conditions. N=3, one-way ANOVA with Šidák’s test, *p < 0.05, **p < 0.01, ***p < 0.001. B) Representative Western Blot analysis from 3 replicates of FTH, TfR1, and GPX4 protein expression. GAPDH was used as a loading control. C) Quantification of protein expression levels (i- FTH, ii- TfR1, iii- GPX4) shown in Panel B. Expression was normalized to the GAPDH loading control and is presented as the relative protein expression, showing upregulation of FTH and downregulation of TfR1 as expected as a result from iron-loading of the cells in presence of FAC and decreased expression of GPX4 in ferroptotic condition with iron-loading and RSL3 treatment.

**Table 1: Top ranked Cell Painting Features in HMC3 ferroptosis model**

Table presenting the top ranked predictive features distinguishing treated HMC3 (iron-laden, RSL3 treated or IronRSL3-treated) from healthy control cells. The ranking was done using a Random Forest classifier. The 35 top features for iron vs DMSO, and RSL3 vs DMSO, as well as the 40 top features for IronRSL3 vs DMSO are listed with the corresponding cellular compartment.

**Table 2: OMICs differential analysis results**

Table containing differential analysis results in lipidomics, proteomics and transcriptomics for the three comparisons: IronDMSO_vs_DMSO, IronRSL3_vs_DMSO, and IronRSL3Fer1_vs_IronRSL3 in the relevant tabs (“IronRSL3Fer1” was shortened to “Fer1” in the tabs name). The last tab also presents the ether phospholipid species found amongst the DALs.

## Acknowledgments

The authors would like to thank Daniel Rock, U-Ming Lim and Abdul Saboor Sheikh for technical support and Jason M.Uslaner and Eimear Howley for providing feedback on the manuscript.

## Author contributions

SG conceived the project with contribution from JR. RB designed experiments together with SG and JAM. RB performed most experiments and analysed the data with contributions from ET for collecting and performing QC of the OMIC samples and from DMC for the MitoSox. NT, EC and MC generated the lipidomic and proteomic datasets and performed the initial analysis. CW performed the computational analysis to integrate all the OMIC data. AL performed the analysis of the Cell Painting data. SG and JAM supervised the work. RB, JAM and SG wrote the manuscript with contributions from JR, NT and CW. All the authors read the manuscript.

## Disclosures

All authors are employees of Merck Sharp & Dohme LLC, a subsidiary of Merck & Co., Inc., Rahway, NJ, USA known as MSD outside of the US and Canada. SG are shareholders of Merck & Co., Inc., Rahway, NJ, USA.

